# Mouse Adapted Omicron BA.5 Induces A Fibrotic Lung Disease Phenotype in BALB/c Mice

**DOI:** 10.1101/2025.07.16.665104

**Authors:** John M. Powers, Sarah R. Leist, Naveenchandra Suryadevara, Seth J. Zost, Elad Binshtein, Anfal Abdelgadir, Michael L. Mallory, Caitlin E. Edwards, Kendra L. Gully, Miranda L. Hubbard, Mark R. Zweigart, Alexis B. Bailey, Timothy P. Sheahan, James E. Crowe, Stephanie A. Montgomery, Jack R. Harkema, Ralph S. Baric

## Abstract

Following SARS-CoV-2 Omicron BA.1, subsequent Omicron sub-lineages have continued to emerge, challenging the development of intervention and prevention strategies, including monoclonal antibodies and vaccines. To better understand the pathogenic effects caused by Omicron BA.5 infection, we developed a mouse-adapted virus with overt disease burden in BALB/c mice. Acute disease was characterized by significant weight loss and lung dysfunction following high-dose challenges. In survivor animals that were followed through 107 days post-infection, subpleural fibrosis with associated tertiary lymphoid structures was noted. Serum from these mice demonstrated potent neutralization against BA.5, with substantially reduced neutralization titers against early epidemic, zoonotic, and more recent contemporary XBB.1.5 variants. Intervention with pre-clinical monoclonal antibodies revealed that robust protection from BA.5-induced lung disease was possible after prophylactic administration. Together, this model enables the investigation of therapeutic approaches for both acute and post-acute sequelae of COVID-19.

**Importance:** In order to best combat the evolving landscape of SARS-CoV-2 variants of interest and variants of concern the development of effective small animal models is of critical importance. Herein, we describe the development of a model system in BALB/c mice to study the effects of SARS-CoV-2 BA.5 in both acute and chronic disease manifestations. Intriguingly, we determined that fibrotic lung disease with tertiary lymphoid structures was a prominent feature in the lungs of mice that survived through the acute phase of infection. This is a prominent concern in human patients that survive the initial infection insult. As such, and most critically, the model system presented here provides researchers with an effective pathway in which long COVID manifestations and potential interventions can be studied.

## Introduction

Severe Acute Respiratory Syndrome Coronavirus 2 (SARS-CoV-2) virions contain an approximately 30 kb positive-sense single-stranded viral genome that encodes numerous structural (spike, membrane, envelope, and nucleocapsid), non-structural proteins (nsp1 – 16), and several accessory genes (1, 2). Following its emergence in late 2019, SARS-CoV-2 rapidly evolved into numerous Variants of Interest (VOI) and Variants of Concern (VOC). The Omicron B.1.1.529 VOC (BA.1) was of significant interest because it encoded a substantial number of amino acid changes within the main antigenic site, the spike gene (S), especially within the receptor-binding domain (RBD), and the receptor-binding motif (RBM). BA.5, which emerged in early 2022, is a descendant lineage of BA.1 that contained additional alterations within the RBD/RBM protein sequence. Due to the role of the spike protein (S) in cellular attachment to angiotensin-converting enzyme 2 (ACE2) and subsequent viral entry following its cleavage by transmembrane protease serine 2 (TMPRSS2), the expanded spike variation was associated with enhanced receptor binding, antibody evasion, and reduced efficacy of both natural and vaccine-induced immunity (3). Amino acid mutations in these domains enabled escape from neutralizing antibodies (4–9).

Inconsistent results in mouse models using BA.5 isolates were observed by different research groups regarding the degree of pathogenicity as compared to BA.1 or previous ancestral viruses (10, 11). However, in both cases, disease severity was overall attenuated compared to early VOCs, like Alpha and Delta, likely mitigated by differences in the clinical isolate, inoculum dose, mouse strain, sex, and age of animals (12–14). Consequently, we generated a mouse-adapted BA.5 virus (designated BA.5 MA), and a derivative strain in which the viral ORF7a was replaced with a gene encoding the reporter protein nanoluciferase (nLuc) to be used as an indicator virus in live-virus neutralization assays (BA.5 nLuc) (15–20). BA.5 MA infection in young (14- to 16-week-old) or aged (10- to 12-month-old) female BALB/c mice caused severe diffuse alveolar damage with accompanying mortality and high virus titers in the lungs and nasal turbinates, which was significantly attenuated by prophylactic administration of monoclonal antibodies.

After symptomatic or asymptomatic infection, SARS-CoV-2 disease may progress to post-acute sequelae of SARS-CoV-2 (PASC), a frequent chronic disease syndrome that includes a continuous, relapsing and remitting, or progressive disease state that affects one or more organ systems and lasts for weeks to years in about ∼10% of SARS-CoV-2 survivors (21–23). In the respiratory tract, shortness of breath, cough, persistent fatigue, post-exertional malaise, and diagnosable conditions like interstitial lung disease and hypoxemia are among the most frequent chronic disease phenotypes. Omicron-related infections are reported to cause less frequent PASC disease in humans as compared to ancestral VOCs (24). After acute BA.5 MA infection, we followed surviving animals through 107 days post-infection (dpi). Mice developed an organizing pneumonia, tertiary lymphoid structures, and pulmonary lung fibrosis as early as 15 dpi, which persisted throughout the study. In contrast, infection with BA.2 MA caused minimal, if any, long COVID disease. Serum from the chronically infected cohort efficiently neutralized BA.5 but was notably less efficient against ancestral or contemporary isolates, suggesting that SARS-CoV-2 contemporary vaccines or natural infections may elicit poorly protective responses to SARS-CoV-2 or more distant Sarbecovirus zoonotic precursor strains. Although speculative, these data suggest that a SARS-like CoV emergence event from zoonotic reservoirs could occur in ∼30+ years or more, as age-related attrition shifts human population level immunity from early pandemic strain exposures toward more highly evolved and antigenically distant contemporary SARS-CoV-2 VOCs. Thus, in the future, VOC-infected newborns/children aged 5 to 10+ years old may exhibit little if any immunity against the zoonotic pool of circulating early pandemic strains.

## Results

### Recovery of Omicron BA.5 Recombinant Viruses

As described in the methods, we recovered a mouse-adapted strain of the BA.5 subvariant isolate (BA.5 MA) and a derivative that expressed nanoluciferase in place of ORF7a (BA.5 nLuc) (15–17). Genome schematics of the infectious clone constructs for the BA.5 MA and nLuc versions are shown in **Supplemental Figure 1**. To assess replication competency, the growth of the two BA.5 recombinant viruses was compared to that of the ancestral strains SARS-CoV-2 MA10 and D614G nLuc, at an MOI of 0.001. In both cases, earlier ancestral strains grew to higher maximal titers than the BA.5 MA and BA.5 nLuc recombinant viruses over the course of 72 hours (**Fig. 1**).

**Figure 1.**
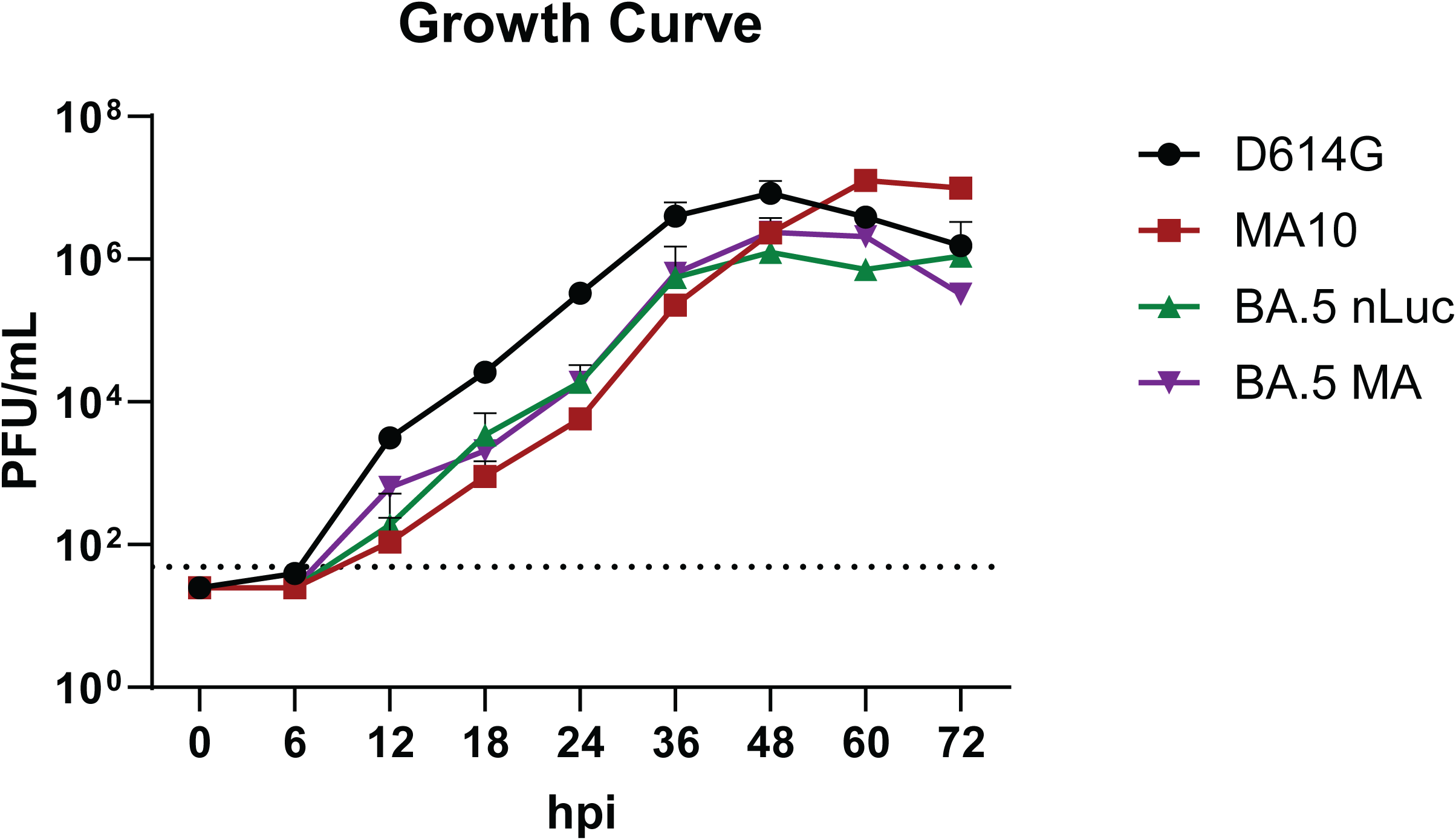
Growth Curve Analysis Identifies Replication Competency of BA.5 MA and nLuc. Growth dynamics for the recombinant BA.5 MA and BA.5 nLuc viruses were compared to ancestral counterparts (D614G nLuc and MA10) to assess for replication competency and equivalent or attenuated replication. Maximal virus titers were reached between 48- and 60-hours post-infection with an MOI of 0.001. Both parental viruses achieved maximal titers ∼10-fold higher than those of the recombinant BA.5 viruses.

### Pathological Features of BA.5 MA Infection in Young Mice

To evaluate BA.5 MA pathogenic outcomes, we infected 14- to 16-week-old adult female BALB/c mice intranasally with different doses, 10^4^ or 10^5^ plaque-forming units (PFU), evaluating dose-dependent disease outcomes. While both cohorts experienced weight loss, the low-dose cohort only experienced a 10.7% group mean weight loss at 4 dpi. In contrast, the high dose cohort continued to lose weight throughout the study, eventually exceeding 20% of starting group mean weights (**Fig. 2A**). Gross lung discoloration (GLD) score is a semiquantitative measure of acute lung damage associated with emerging CoV replication, indicative of edema and diffuse alveolar damage (17, 25). Similar to weight loss, GLD scores were significantly increased in the high-dose cohort as compared to the low-dose group (**Fig. 2B**). Similarly, mortality was only observed in the high-dose group (**Fig. 2C**). Regardless of inoculum dosage, the levels of viral replication within the lungs and nasal turbinates showed no significant difference between the two cohorts (**Fig. 2D & E**). Acute lung injury (**Fig. 2F**) and diffuse alveolar damage (**Fig. 2G**) were assessed using histological scoring schema that have been utilized for multiple emerging CoV (17, 25). Both scoring metrics (**Fig. 2F & G**) revealed that acute lung injury was significantly elevated regardless of virus dose which was evident in photomicrographs of representative mock-infected (PBS inoculation) (**Fig. 2H**), 10^4^ PFU (**Fig. 2I**), and 10^5^ PFU (**Fig. 2J**) infected animals sacrificed at 4 dpi.

**Figure 2.**
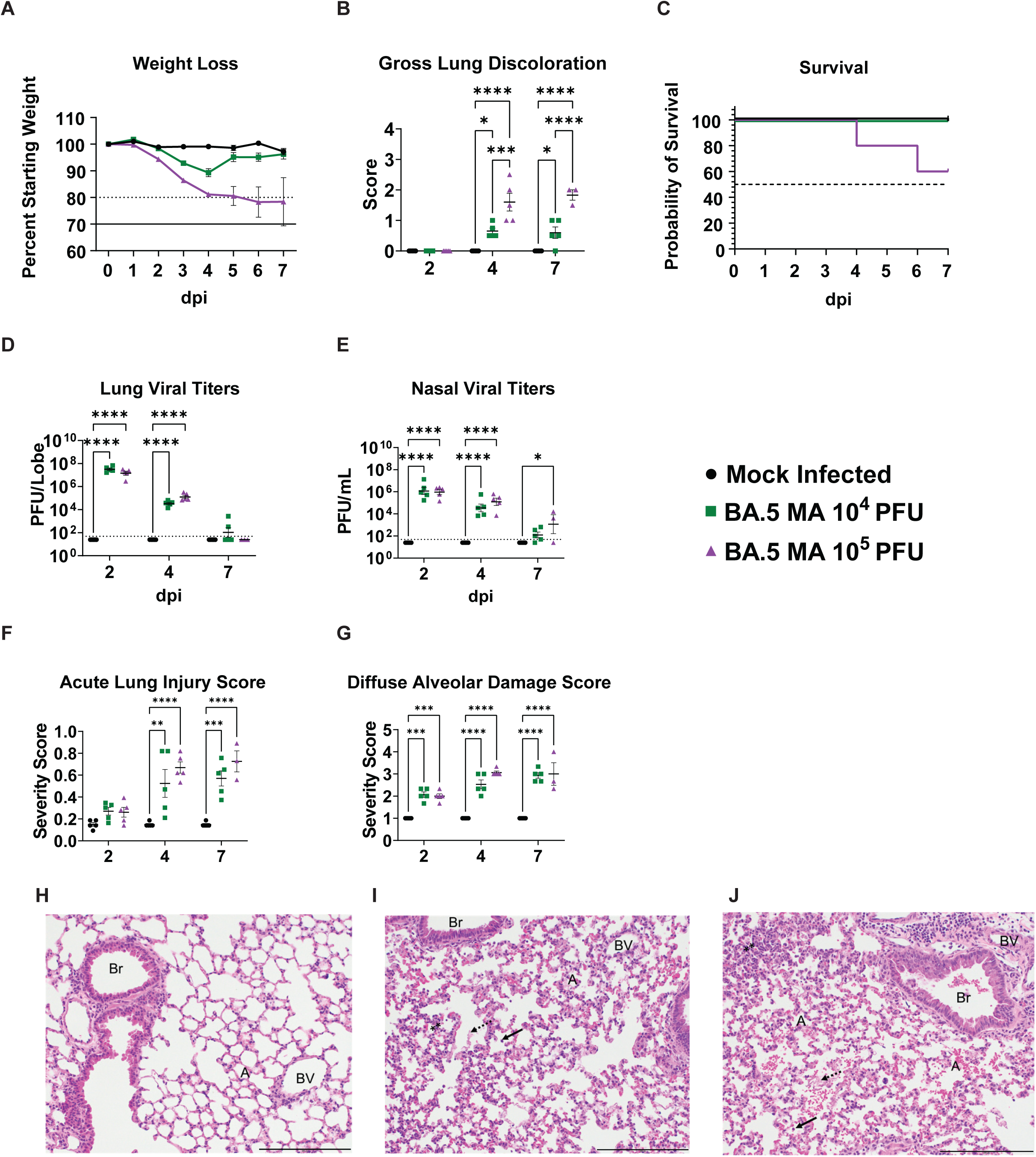
Dose-Dependent Pathogenicity in Young Mice Following BA.5 MA Challenge. Pathogenicity of the BA.5 MA virus was assessed in 14- to 16-week-old female BALB/c mice to determine their applicability as a model system for pathogenicity studies. (**A**) Weight loss was tracked daily for PBS control cohorts, as well as 10^4^ and 10^5^ PFU BA.5-MA-infected mice. Dashed line represents 20% weight loss and solid line represents 30% weight loss, a humane euthanasia criterion. (**B**) Gross lung discoloration was evaluated and scored at the indicated timepoints of 2, 4, and 7 dpi. (**C**) Survival analysis of mice in the 7 dpi cohorts identified only a dose of 10^5^ PFU resulted in mortality. (**D & E**) Virus titers were determined for replication-competent virus by plaque assay following homogenization of lung or nasal turbinate tissues. Dashed line represents limit of detection. (**F & G**) Lung damage was evaluated by two metrics, either Matute-Bello acute lung injury scoring in panel **F** or by diffuse alveolar damage scores in panel **G**. (**H – J**) Representative histopathological sections of lungs from PBS (**H**), 10^4^ PFU BA.5 MA (**I**), or 10^5^ PFU BA.5 MA infected mice (**J**) at 4 days post-infection are shown (A, alveoli; Br, bronchiole; BV, blood vessel; stippled arrows, proteinaceous debris; ⁎⁎, hypercellular alveoli; solid arrow, presence of infiltrating cells in alveolar spaces). Scale bar represents 200 μm.

### Pathological Features of Infection in Aged Mice

Viral pathogenesis was then evaluated in 10- to 12-month-old mice infected with 10^4^ or 10^5^ PFU of BA.5 MA. Both high- and low-dose cohorts rapidly lost weight approaching 80% of starting weight by 4 dpi (**Fig. 3A**). GLD scores were elevated in the high dose cohort as compared to the low dose group (**Fig. 3B**). Although trends in body weight loss were similar up to 5 dpi, high dose infection was uniformly lethal (**Fig. 3C**) while the majority of low dose infected animals survived out to 15 dpi. As observed in the younger mice above, lung and nasal titers between the infected groups did not differ significantly, regardless of dose (**Fig. 3D & E**). As expected, infection was associated with increases in the histopathological measures of acute lung injury (**Fig. 3F**) and diffuse alveolar damage (**Fig. 3G**) during acute infection. Counterintuitively, the low dose group had elevated histological scores as compared to the high dose group, but this trend was likely driven by survivor bias where the high dose infected animals that survived to later times post infection likely had diminished disease severity as compared to those that perished at early times post infection. Representative images are shown at 4 dpi for mice inoculated with PBS, 10^4^ PFU, or 10^5^ PFU of BA.5 MA (**Fig. 3H – J**). At later times post infection (7, 15 dpi) we evaluated tissue sections for the prevalence of pro-fibrotic regions (**Fig. 3K**). Like before, only the low dose group had elevated fibrosis scores over mock infected animals and importantly no high dose animals survived to 15 dpi thus could not be included in this analysis.

**Figure 3.**
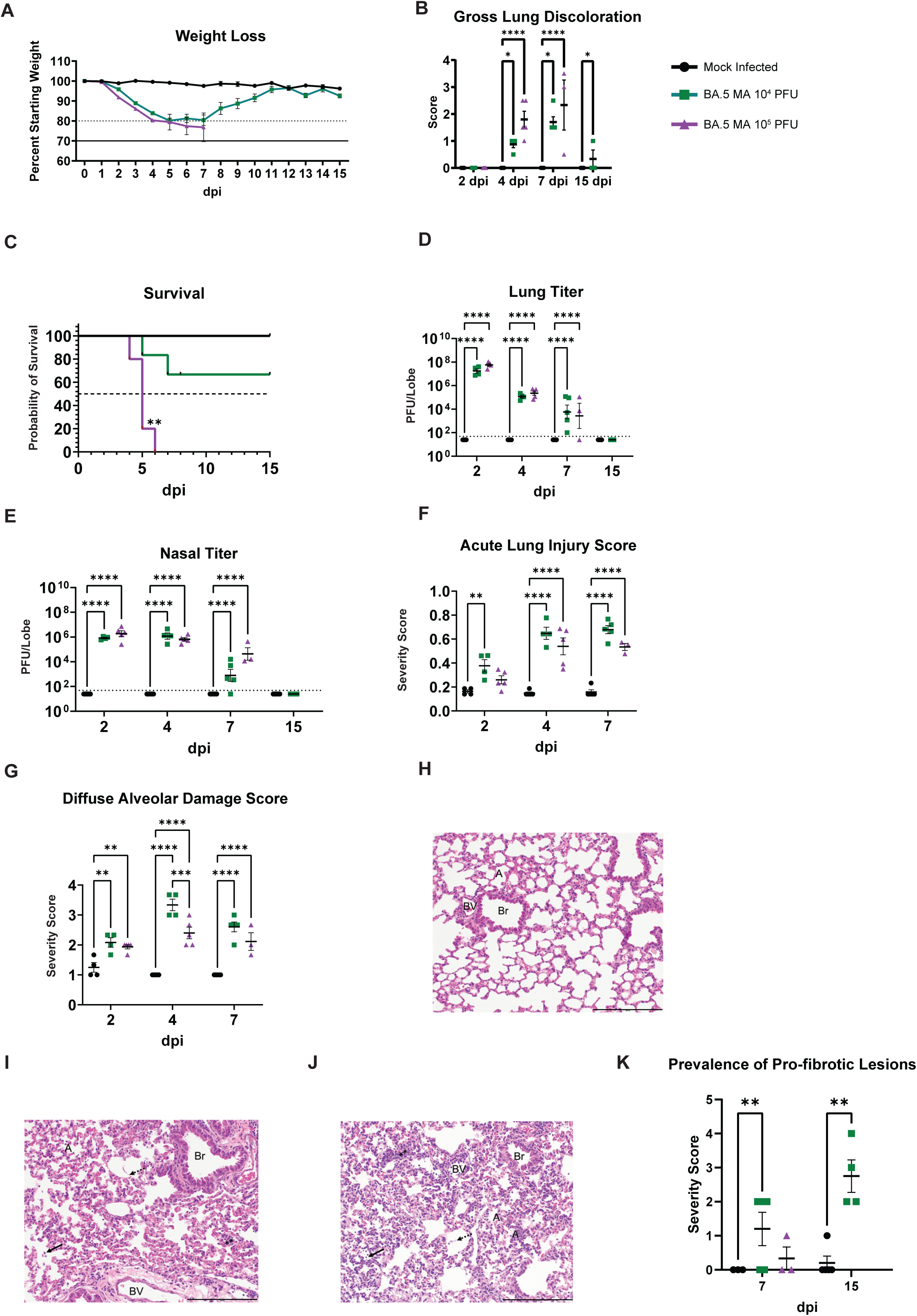
BA.5 MA induces Pro-Fibrotic Lesions in Aged Mice Following Acute Phase Disease Resolution. 10- to 12-month-old female BALB/c mice were used to evaluate the pathogenicity after receiving a dose of either 10^4^ or 10^5^ PFU of BA.5 MA. (**A**) Weight loss was recorded daily over the course of the study. The dashed line represents 20% weight loss, and the solid line represents 30% weight loss, a humane euthanasia criterion. (**B**) Gross lung discoloration was recorded at each harvest timepoint. (**C**) Mortality of a representative cohort of five mice per group that were followed until the end of the study at 15 dpi. (**D & E**) Plaque assays were conducted on lung and nasal turbinate tissues to determine replication-competent virus burden. Dashed lines represent limit of detection. (**F & G)** Histopathological examination was conducted on fixed lung sections to determine lung damage as a result of challenge. (**H – J**) Representative images are shown at 4 dpi for mice inoculated with PBS, 10^4^ PFU, or 10^5^ PFU (A, alveoli; Br, bronchiole; BV, blood vessel; stippled arrows, proteinaceous debris; ⁎⁎, hypercellular alveoli; solid arrow, presence of infiltrating cells in alveolar spaces). Scale bar represents 200 μm. (**K**) Prevalence of pro-fibrotic lesions was scored for lungs from subjects following Picrosirius red staining (0 = none; 1 = <5% of parenchyma; 2 = 6 to 10%; 3 = 11 to 50%; 4 = 51 to 95%, 5 = >95%).

An age matched cohort of mice was inoculated with a BA.2 MA virus that expressed the BA.2 spike gene and MA10 backbone mutations, as described previously (17). Mice were challenged with a single dose of 10^5^ PFU, and the relative disease state was compared between the two challenge viruses. Compared to the BA.5 MA-infected cohort, BA.2 MA-infected mice showed a maximal weight loss of 3.7% (**Fig. S2A**), with weight loss profiles closely matching those of the mock-infected control cohort. While weight loss was not a feature of disease, viral replication in both the lung and nasal turbinates reached comparable levels to those seen in the BA.5 MA-infected mice, demonstrating the impact of spike protein variation on disease progression and severity (**Fig. S2B**). Like weight loss data, lung discoloration was not a feature of disease, with minimal observable scoring noted at 4 dpi (**Fig. S2C**).

### Monoclonal Antibody Treatment Abrogates BA.5 MA-Inflicted Disease

From a selected panel of representative SARS-CoV-2 reactive human monoclonal antibodies (mAbs), we identified two that showed robust neutralization titers against BA.5 nLuc (**Fig. 4A**). Two mAbs (COV2-3605 and COV2-3678) were compared against a recombinant version of a neutralizing SARS-CoV-2 monoclonal antibody (rLY-CoV1404) that served as a positive control, and an isotype-matched recombinant version of a human mAb antibody recognizing the unrelated dengue virus envelope protein (rDENV-2D22) that served as a negative control. In addition, a mock treatment group received an equivalent volume of PBS. In a neutralization assay against BA.5 nLuc, all three SARS-CoV-2 antibodies potently neutralized the virus, with IC_50_ titers of 9.7 ng/mL for COV2-3605, 51.7 ng/mL for COV2-3678, and 3.2 ng/mL for rLY-CoV1404 (**Fig. 4A**). Both, COV2-3605 and COV2-3678, are encoded by the human antibody variable gene segment *IGHV3-53*. Many *IGHV3-53*- and *IGHV3-66*-encoded mAbs constitute a public clonotype, and this class of antibodies has been recurrently isolated from human subjects following SARS-CoV-2 infection or vaccination (26–29). Negative-stain electron microscopy revealed COV2-3605 Fab and COV2-3678 Fab bound to RBDs in the open conformation of the spike, consistent with the fact that members of this public clonotype target the semi-cryptic class I antigenic site (**Fig. 4B**) (30). Negative-stain electron microscopy data collection statistics are provided in **Supplemental table 1**.

**Figure 4.**
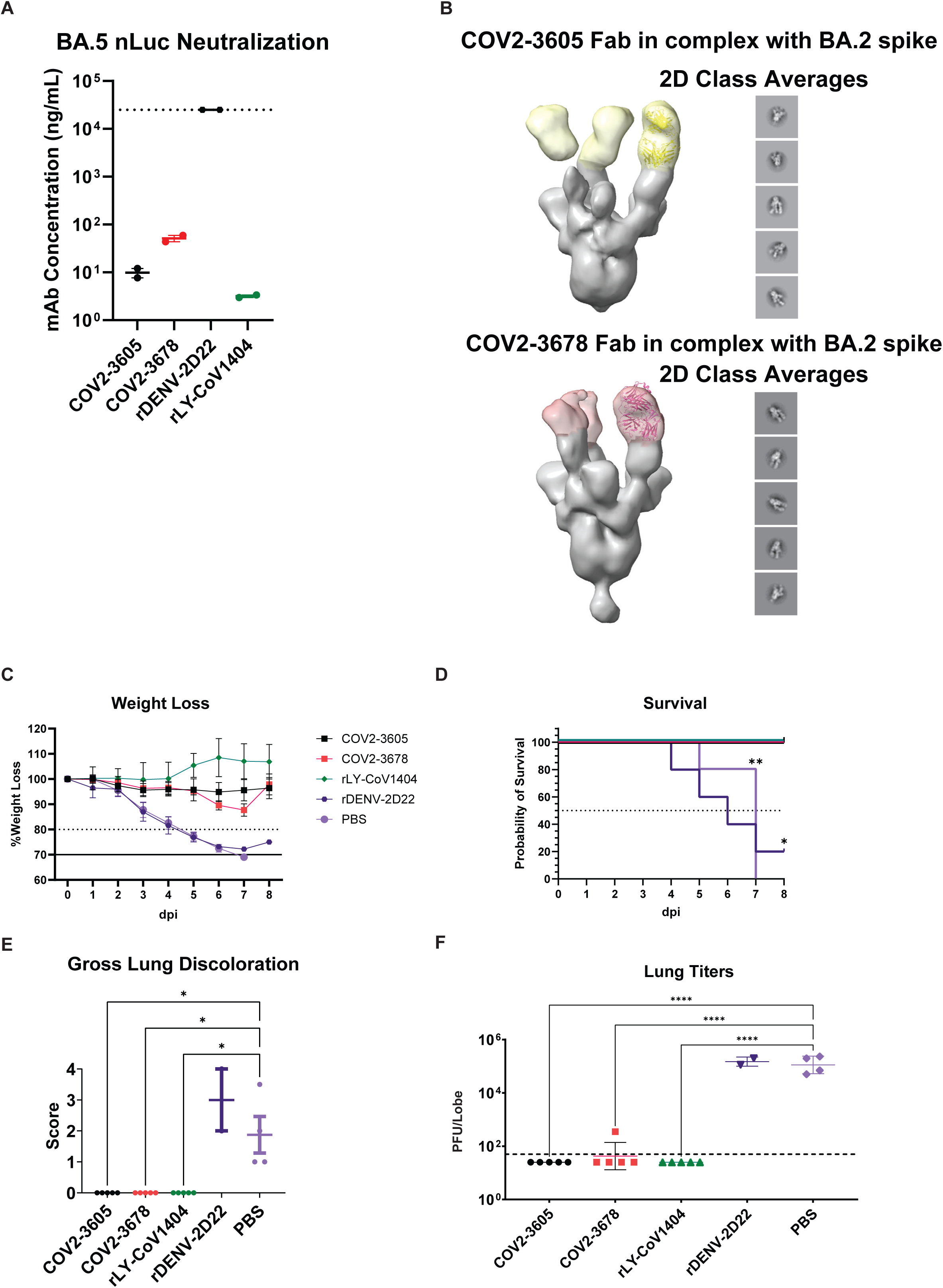
Candidate Monoclonal Antibodies Abrogate Disease Pathology. 10- to 12-month old female mice were dosed prophylactically with 200 μg of mAb or PBS intraperitoneally 12 hours prior to challenge. (**A**) Neutralization potency of mAbs was assessed against BA.5 nLuc. (**B**) Negative-stain EM of COV2-3605 and COV2-3678 Fabs in complex with BA.2 S protein shows these mAbs recognize the RBD in the up conformation and likely bind the Class I antigenic site. (**C**) Weight loss was recorded daily over the course of the study. The dashed line represents 20% weight loss, while the solid line represents 30% weight loss, a humane euthanasia criterion. (**D**) Cohort survival was evaluated for a subset of mice (n=5) that was followed for the entire duration of the study. (**E**) Gross lung discoloration was observed at the time of mouse sacrifice. (**F**) Lung titers were determined via plaque assay to assess the quantity of replication-competent virus in lung tissue taken from mice sacrificed at 4 dpi. The dashed line represents the limit of detection.

To assess the prophylactic efficacy of mAbs, 10- to 12-month-old female BALB/c mice were treated with antibodies via intraperitoneal injection of 200 μg of mAb, and 12 hours later, mice were inoculated with a lethal dose of 10^5^ PFU of BA.5 MA. Mice treated with COV2-3605 or rLY-CoV1404 showed minimal or no weight loss. In contrast, ∼10% transient weight loss was observed at 5 and 6 dpi for the COV2-3678-treated cohort (**Fig. 4C**). Importantly, negative control group mice (i.e. PBS or isotype-matched control mAb rDENV-2D22) showed rapid weight loss through 7 dpi, with all animals succumbing to infection or reaching humane endpoints for euthanasia by 7 – 8 dpi (**Fig. 4D**). Similarly, only negative control group animals had elevated GLD scores (**Fig. 4E**). Minimal breakthrough infection was noted in the COV2-3678-treated mice, with one of five mice exhibiting low replicating virus at 4 dpi (**Fig. 4F**). In contrast, no live virus was detected in the COV2-3605 and rLY-CoV1404 mAb-treated cohorts. Viral titers in mAb-treated cohorts were significantly reduced compared to PBS-treated cohorts, which had titers approaching 10^5^ PFU/lobe (p < 0.0001).

### Chronic Model of BA.5 MA Lung Disease

To determine the potential for BA.5 MA and BA.2 MA to cause PASC disease phenotypes in the lung, cohorts of 10- to 12-month-old female BALB/c mice were inoculated with 10^4^ PFU or mock-treated and followed through 107 dpi (BA.5 MA) or 120 dpi (BA.2 MA). Cohorts of five animals were sacrificed at 15, 30, 60, 90, or 107/120 dpi. Weight loss, mortality, gross pathology, and viral titers were evaluated over time.

BA.5 MA infected mice experienced rapid weight loss with mean reductions of 17.4% ± 7.6% by 7 dpi, and recovery of body weight was gradual, finally achieving similar weights to mock-infected controls by 30 dpi (**Fig. 5A**). Congruent with **Figure 3**, BA.5 MA infection diminished survival during the acute phase with only 60% of the initial cohort of infected mice surviving beyond 9 dpi (**Fig. 5B**). GLD scores were low but measurable at 15 dpi but waned over time (**Fig. 5C**). Neither replication competent virus via plaque assay nor viral RNA via qRT-PCR was detected in lung tissue (data not shown). Additionally, viral nucleocapsid antigen was not detected in tissue sections of lung, liver, kidney, or spleen at any of the times assessed (data not shown).

**Figure 5.**
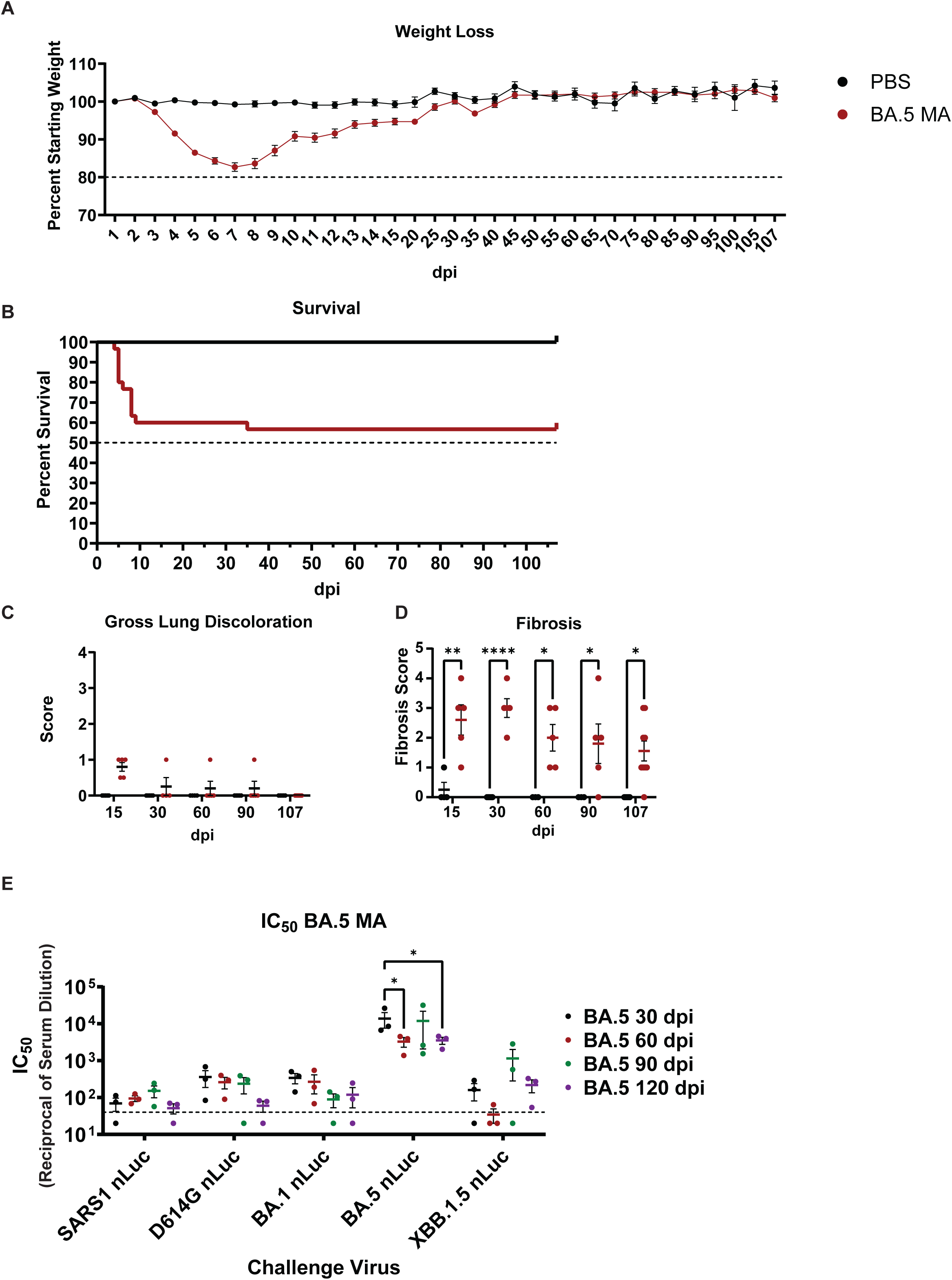
BA.5 MA Induces Chronic Lung Disease Phenotypes and Induces Persistent Homotypic Neutralizing Antibodies. 10- to 12-month-old female mice were inoculated with 10^4^ PFU of BA.5 MA and followed for 107 days. (**A**) Weight loss was tracked daily over the acute phase of the infection (15 days), after which it was recorded every five days. (**B**) Survival was recorded for mice from the 107 dpi cohort. (**C**) Lung discoloration was evaluated at the indicated timepoints. (**D**) Histopathological examination was performed on Picrosirius-red-stained lung sections to determine the presence of fibrotic lesions in the lung parenchyma. (**E**) Live-virus neutralization assays were performed on blood serum collected from mice at the indicated collection timepoints. Sera were assayed against SARS-CoV or SARS-CoV-2 viruses expressing the D614G spike or Omicron BA.1, BA.5, and XBB.1.5 S proteins. Dashed line represents limit of detection.

As with the acute-phase infection and in contrast to BA.5 MA, BA.2 long-term study mice displayed minimal to no weight loss (**Fig. S2A**), no mortality, no apparent GLD scores (not shown), and minimal lung fibrosis scores (0.4) of the total parenchyma by Picrosirius red staining (0 = none; 1 = <5%; 2 = 6 to 10%; 3 = 11 to 50%; 4 = 51 to 95%, 5 = >95%) (**Fig. S2D**). In contrast, BA.5 MA group mean scores averaged between ∼2 to 3 over time, indicating fibrotic lesions involving 6% to 50% of the lung parenchyma (**Fig. 5D**). By day 107, about 20% of the BA.5 MA virus infected mice had resolved the most prominent long COVID lesions. While scores varied between the individual animals within a given cohort and timepoint, the data show the range in which chronic disease can manifest and present. Representative images of H&E and Picrosirius red (PSR) stained slides are shown for mock-infected and BA.5 MA infected animals at 30 and 107 dpi (**Fig. S3**). Pulmonary histopathology was not observed in mock-infected mice. In contrast, BA.5 mice at 30 or 107 dpi exhibited subpleural chronic alveolitis with scattered tertiary lymphoid structures and areas of PSR-stained interstitial fibrosis in the alveolar parenchyma.

### Cross-Neutralization Antibody Titers After Natural Infection

We next measured the magnitude and durability of the serologic neutralizing response in BA.5 MA challenged mice using homotypic (BA.5 nLuc), or heterotypic (SARS-CoV-1 nLuc, SARS-CoV-2 D614G nLuc, SARS-CoV-2 BA.1 nLuc, and SARS-CoV-2 XBB.1.5 nLuc) Sarbecoviruses (20). Using live nLuc reporter viruses, we observed peak homotypic neutralization titers of >1/10,000 against BA.5 by 30 dpi, however, titers waned quickly and were significantly reduced by 5- to 10-fold through 107 dpi (p < 0.05) (**Fig. 5E**). In contrast, levels of heterotypic neutralization titers were greatly reduced when compared to former ancestral strains (*e.g*., SARS-CoV-2 D614G, SARS-CoV-2 BA.1), contemporary strains (*e.g*., SARS-CoV-2 XBB.1.5), and early 2003 epidemic isolates (SARS-CoV-1) (**Fig. 5E**). While infection leads to a durable homotypic neutralizing antibody response through ∼4 months, neutralization potency waned rapidly over time, especially against ancestral and zoonotic strains. Similar findings were noted with serum from day 120 BA.2 MA neutralizing antibody titers (**Fig. S2E**).

## Discussion

SARS-CoV-2, the etiological agent responsible for the COVID-19 pandemic, has caused significant global morbidity and mortality, with excess mortality estimates approaching 20-25 million or more (31). Although the explosive epidemic phase of the pandemic has transitioned into an endemic phase, with an estimated 32,000 to 50,000 deaths from October 2024 through June 2025, mortality remains concentrated among the elderly and certain immunocompromised populations (32–34). Importantly, in most populations, acute disease severity of the Omicron lineage strains is reduced compared to that caused by early ancestral VOCs, which is also reflected in animal models. Despite achieving similar titers in the mouse, our data show that S protein sequence variation between Omicron strains in an isogenic backbone is a significant determinant of long COVID pulmonary disease traits. Reduced long COVID disease severity likely reflects the role of S gene variation in virulence and pathogenesis, and S variation likely contributes to increased long COVID disease rates after secondary and tertiary infections in humans (35). Importantly, antigenic variation, coupled with the higher transmission efficiency of Omicron lineage strains, has caused significant acute and chronic healthcare burdens worldwide (36, 37).

Both early and contemporary SARS-CoV-2 VOC infections cause chronic multiorgan system-level disease phenotypes, which have been termed long COVID or PASC. Long COVID disease symptoms persist for at least 3 months after resolution of acute infection, oftentimes longer in the brain and lung (38, 39). While repeat infection is associated with enhanced and/or reactivated long COVID disease phenotypes and vaccines reduce the prevalence of long COVID, it remains uncertain as to whether this trend will continue as new VOCs emerge, and whether pre-existing long COVID conditions will alter the severity of acute and chronic disease risk for new VOCs or even other respiratory virus infections. The development of early and contemporary SARS-CoV-2 acute and chronic disease models, like MA10, BA.2 MA, and BA.5 MA, provides platforms to address these and related questions in future studies, such as the impact of repeated lung injury from SARS-CoV-2 VOC infections on long-term disease phenotypes. While the mechanisms and complex traits that predispose some individuals to acute COVID-19 pathogenesis are encoded by many common loci and underlying genes in humans and rodent genetic reference populations (40–45), the host genetic drivers of long COVID disease are confounded by pre-existing conditions, immune status, co-morbidities, and medications, coupled with environmental factors, virus dose/strain, repeat infections, sex, vaccination status, hospitalization status, critical care modalities, and complex case definitions of ∼200+ long COVID disease phenotypes (46). Chronic PASC animal models, like the one we describe for BA.5 MA, are needed to understand disease mechanisms, identify genetic susceptibility loci and candidate disease-driving genes, and pathways that inform countermeasure performance (38, 47, 48). For example, although several human studies have suggested frequent long-term viral persistence as a principal driver of PASC, only about 10-20% of cases demonstrate prolonged persistent viral RNA, and a 15-day treatment with nirmatrelvir + ritonavir (Paxlovid) was safe, but did not demonstrate clinical improvement (46, 49). Our data, and those of others, support another potential model that drives long COVID disease in which aberrant epithelial-immune cell interactions and lung repair defects are major drivers of chronic disease (50).

In pre-immune populations, Omicron variants, like BA.5, usually cause less severe acute symptoms and infections, and patients are generally at lower risk for post– COVID–19 conditions than after infection with an ancestral VOC (10, 51–54). In other areas across the globe, about 11.8% of patients with omicron-dominant infections developed post–COVID-19 conditions, most often in females (55). Moreover, the considerable S protein antigenic variation in Omicron-related strains has eroded the performance of natural infection and vaccine protective immunity. To address these issues, we developed a BA.5 MA dose-dependent acute lung injury model that captures age-related disease susceptibility and can progress to fatal outcomes, and a significant proportion of survivors (18/20) developed a chronic organizing pneumonia with fibrosis through day 107. After day 90, two of 10 animals appeared to resolve nearly all of the long COVID disease phenotypes, suggesting effective reparative mechanisms operating over time. Although less pathogenic, the model captures many of the Wuhan MA10 pathogenesis phenotypes, including early acute respiratory disease syndrome phenotypes that progress to chronic organizing pneumonia with fibrosis in young and aged animals after acute and chronic infection, as previously reported by our group (17, 56). The model reflects disease patterns noted in human cohorts (32, 57). These data in mice are consistent with findings in human studies that identify disease severity and age as important risk factors for chronic PASC conditions (58, 59).

Due to the clear evidence of disease during the acute phase of infection, the BA.5 model serves as a reliable and robust platform for evaluating countermeasure performance. Monoclonal antibody therapies have significantly reduced the amount of life-threatening disease in human populations (60, 61). With the emergence of the Omicron lineage of SARS-CoV-2, significant declines in the performance of previously approved mAbs were observed (62–64). The accrual of mutations throughout the S protein, especially those changes in the RBD and RBM of the S protein, enables decreased susceptibility to the authorized antibodies (65). We evaluated the capacity of two mAbs (COV2-3605 and COV2-3678) to limit disease burden when provided prophylactically before challenge. These antibodies were isolated from an individual following a BA.1 breakthrough infection, target the RBM of the SARS-CoV-2 RBD, and are encoded by antibody variable gene *IGHV3-53*, making them members of a previously described public clonotype using *IGHV3-53/IGHV3-66* genes. Consistent with their potent neutralization of the BA.5 variant (**Fig. 4A**), both antibodies significantly attenuated disease progression after BA.5 MA infection (**Fig. 4C – F**). Importantly, the class I antigenic site targeted by members of this public clonotype continues to be a major target of human immunity and has accumulated further antigenic substitutions (66). However, some mAbs with this gene usage have been described that show extraordinary breadth of reactivity and resilience to escape (67).

Like SARS-CoV-2 MA10 (based on the Wuhan virus strain), the BA.5 MA virus causes diffuse alveolar damage and severe acute lung injury through 7 dpi. Survivors through 107 dpi revealed significant chronic long COVID clinical and pathologic phenotypes with lung fibrosis, as determined by picrosirius red staining, noted as early as 15 dpi (**Fig. 5D**). As the low-dose BA.5 MA model reliably instilled an organizing pneumonia with tertiary lymphoid structures and fibrotic lesions in the lung through 107 dpi with minimal mortality, future interventional approaches could be evaluated to determine its utility in evaluating the efficacy of vaccines, therapeutics, and small molecule inhibitors to abrogate disease as described previously by our group (16). In contrast, when we compare the results of BA.5 MA to those observed in the BA.2 MA infection cohort, we recognized significant differences in the overall acute and chronic disease outcomes and trajectory for the mice. While these viruses are closely related and share a number of mutations in the S protein, the genome backbone sequences are isogenic, indicating that the observed phenotype is S-driven, despite comparable viral titers in the lung (68). Notably, our group has previously demonstrated that a single amino acid polymorphism in S protein can drive widely disparate disease outcomes in mice infected with XBB.1 MA versus XBB.1.5 MA (20).

Previous studies have revealed that COVID-19 humoral immunity acquired from natural infection is highly variable in magnitude and half-life as compared to vaccine-elicited immunity [38-40]. In general, more severe infections oftentimes elicit higher and more durable neutralizing responses as compared to mild infections in humans, in a host-dependent manner (69–71). To address this question, we evaluated the longevity and breadth of neutralizing antibodies elicited after BA.5 MA and BA.2 MA infection. Using homotypic or heterotypic virus strains, we found that natural infection stimulates strong and durable homotypic neutralizing antibodies through 107 dpi, with only ∼10-fold waning responses. However, neutralization breadth was limited especially against heterotypic ancestral (*e.g*., SARS-CoV-2 D614G, Omicron BA.1) strains, Sarbecovirus clade variants like the 2003 SARS-CoV epidemic strain, and to a lesser extent the more contemporary SARS-CoV-2 XBB1.5 VOC strain (**Fig. 5E**). Among related, contemporary human coronavirus subgenus pairs (group 2a: HCoV OC43; HCoV HKU1; group 1b: HCoV 229E; HCoV NL63), these pairs have been predicted to have emerged centuries apart from unique zoonotic introductions, supporting the hypothesis that sufficient antigenic variation exists in the zoonotic pool to allow for independent CoV re-introduction events in the future (72, 73). While early isolates of the contemporary human CoV strains are not available, the SARS-CoV-2 pandemic provides a critical model to study coronavirus evolution/time, especially in relation to contemporary and zoonotic isolates and future VOCs. After natural infection, BA.2 and BA.5 spikes have antigenically evolved away from early pandemic isolates and the zoonotic pool. So far, human herd immunity has selected for novel antigenic repertoires that radiate away from the earliest SARS-CoV-2 strains and zoonotic reservoirs (74). If these trends continue, human immune neutralizing antibody repertoires targeting the ancestral SARS-CoV-2 will diminish over time because of age-related attrition in human populations. Consequently, in ensuing years and decades, the immune repertoires of newborns who have imprinted on future contemporary “human evolved” SARS-CoV-2 VOCs will govern population-level immunity. This process may occur relatively quickly, as roughly 30 or 50% of the global human population is under 15 or 30 years of age in 2024, respectively (75). As seen with other human Alpha and Beta coronaviruses, our data suggests the potential for a second SARS-CoV-2 re-emergence event in >30+ years, with this likelihood increasing over time as much of the human population retains little cross-protective immunity against distant strains in the zoonotic pool (**Fig. S4**). Moreover, policy changes that allow for BSL2 culturing of Wuhan SARS-CoV-2 under laboratory conditions in many countries, coupled with the persistence of ancestral SARS-CoV-2 strains like Alpha and Delta variants in deer and perhaps other mammals, would lead to opportunities for the re-emerge of SARS-CoV-2. In the latter examples, variants with potentially unique intra-host evolved S protein mutations could provide a new source of antigenically distinct Sarbecoviruses to threaten human populations (**Fig. S4**)(72, 74, 76–79). Disease risk, however, would likely be somewhat mitigated by high levels of circulating T cells and perhaps Fc-mediated protective immunity, unless countered by virulence-enhancing determinants (80–85).

In summary, we show that Omicron-related strains can elicit an acute and chronic pathogenic outcome in BALB/c mice, including progression to long COVID disease in the lungs of survivors. The model can be tuned for disease severity based on initial virus challenge dose and age of animals, resulting in more overt disease symptoms at higher doses or advanced age, although replication fitness was near equivalent regardless of inoculum dosage. Most critically, the model provides the ability to study either acute or chronic lung disease phenotypes, providing a platform in which disease mitigators can be evaluated. A weakness of the study is the focus on disease patterns in female mice, as studies with MA10 have revealed more significant virulent acute and chronic disease phenotypes in the lungs of males (86). Future studies also provide opportunities to evaluate the BA.5 MA infection potential to elicit long COVID disease phenotypes in the brain and olfactory epithelium, as have been described with ancestral SARS-CoV-2 strains (87, 88).

## Materials and Methods

### Virus and Cells

Using reverse genetics, we recovered the BA.2 and BA.5 wildtype spike gene (S) sequence in the background of previously described mouse-adapted mutations (BA.2 MA and BA.5 MA) (GenBank under accession numbers PV800150 & PV800152) [20,21]. A second set of viruses encoding the BA.2 or BA.5 S protein sequence that expressed nanoluciferase gene in place of ORF7a as previously described (BA.2 nLuc and BA.5 nLuc) (GenBank under accession numbers PV800151 & PV800153) [20, 21]. Infectious virus recovery was performed as previously described (16, 20).

For recombinant protein expression, Expi293F cells (Thermo Fisher Scientific; cat # A1435101) were maintained at 37 °C in 8% CO_2_ in Expi293F Expression Medium (Thermo Fisher Scientific; catalog number A1435102), while ExpiCHO cells were maintained at 37 °C in 8% CO_2_ in ExpiCHO Expression Medium (Thermo Fisher Scientific, cat # A2910001).

### Mice and *in vivo* Infections

Female 14- to 16-week-old or 10- to 12-month-old BALB/c mice were obtained from Envigo (Inotiv) (strain 047). Mice were inoculated intranasally under ketamine/xylazine anesthesia with either 1 x 10^4^ or 1 x 10^5^ PFU BA.5 MA, or 1 x 10^5^ PFU BA.2 MA in 50 μL PBS as indicated, as previously described (16, 17, 20).

### MAb and Antigen Production and Purification

cDNAs encoding mAbs of interest were synthesized (Twist Bioscience) and cloned into an IgG1 monocistronic expression vector (designated as pTwistmCis_G1) and used for production in mammalian cell culture. This vector contains an enhanced 2A sequence and GSG linker that allows for the simultaneous expression of mAb heavy and light chain genes from a single construct upon transfection. For antibody production, we performed transfection of ExpiCHO cell cultures using the Gibco ExpiCHO Expression System as described by the vendor (89). IgG molecules were purified from culture supernatants using HiTrap MabSelect SuRe (Cytiva) columns on a 24-column parallel protein chromatography system (Protein BioSolutions).

To express SARS-CoV-2 S proteins for ELISA binding and electron microscopy (EM) studies, we introduced the mutations of the BA.2 variant into the context of a previously described stabilized S protein construct (VFLIP) (90). In addition to a C-terminal T4 fibritin foldon domain, an 8× His tag, and a TwinStrep tag, this construct contains an inter-protomer disulfide bond, a shorter glycine-serine-rich linker between the S1 and S2 domains, and five proline substitutions relative to the native SARS-CoV-2 S sequence. Plasmid encoding the BA.2_VFLIP antigen was transiently transfected into Expi293F cells, and culture supernatants were collected 4 to 5 days following transfection. After clarification by centrifugation and the addition of BioLock (IBA LifeSciences), antigen was purified using affinity chromatography with StrepTrap XT columns.

### Electron Microscopy Sample Preparation

Electron microscopy imaging was performed with COV2-BA2 spike protein in complex with either COV2-3605 or COV2-3678. A recombinant form of the COV2-BA.2 spike was expressed and purified by affinity. Fabs were generated from purified, recombinantly expressed mAbs via enzymatic digestion using a FabALACTICA kit (Genovis, cat # A2-AFK-005). Antigen-Fab complexes were generated by incubating BA.2 S_VFLIP antigen with COV2-3605 Fab or COV2-3678 Fab in a 1:4 (antigen:Fab) molar ratio for 2 hours at room temperature.

### Negative-stain Grid preparation, Imaging and Processing

3 µL of the complex sample at ∼10 µg/mL was applied to a glow-discharged grid with continuous carbon film on 400 square mesh copper electron microscopy grids (Electron Microscopy Sciences). Grids were stained with 2% uranylformate (91). Images were recorded on a Gatan US4000 4k’ 4k CCD camera using an FEI TF20 (TFS) transmission electron microscope operated at 200 keV and controlled with SerialEM (92). All images were taken at 50,000 magnification with a pixel size of 2.18 Å/pixel in low-dose mode at a defocus of 1.5 to 1.8 μm. The total dose for the micrographs was ∼33 e/Å2. Image processing was performed using the cryoSPARC software package (93). Images were imported, CTF-estimated and particles were picked automatically.

The particles were extracted with a box size of 256 pix and binned to 128 pix (4.36 Å/pixel), and multiple rounds of 2D class averages were performed to achieve clean datasets. The final dataset was used to generate an initial 3D volume, and the volume was refined for a final map at the resolution of ∼18 Å. Fab Model docking to the EM map was done in Chimera. PDB: 12E8 was used for the Fab. ChimeraX software was used to make all the figures (94). Data collection statistics are provided in **Supplemental table 1**.

### Assessment of MAbs *in vivo*

Antibody studies were conducted with mice treated prophylactically (12 hours prior to infection) with 200 μg of the indicated mAbs or isotype-matched IgG controls. Virus-challenged mice were inoculated with BA.5 MA at 1 x 10^5^ PFU intranasally. A mock-infected control cohort received an equivalent volume of phosphate-buffered saline intranasally.

### Nanoluciferase-Based Neutralization Assays

Moderate-throughput nanoluciferase assays in a 96-well format were conducted as previously described (20).

### Histopathology and Antigen Staining

Following harvest, mouse lungs were fixed for ≥7 days in 10% phosphate-buffered formalin at 4 °C and prepared for histological examination and scoring as previously described (16, 20).

### Statistical Analysis

All statistical analyses were performed using GraphPad Prism 10. Statistical significance was determined by two-way ANOVA with Tukey’s multiple comparison test for weight loss, Kruskal-Wallis non-parametric test with Dunn’s correction for lung discoloration, and one-way ANOVA with Tukey’s multiple comparison test for tissue titers. * = p < 0.05, ** = p < 0.01, *** = p < 0.001, **** = p < 0.0001. All cohorts started with five mice per harvest timepoint at the time of infection. Error bars represent standard error of the mean.

### Ethics and Containment Procedures

Work with recombinant viruses was approved by the University of North Carolina at Chapel Hill Institutional Review Board under Schedule G 78684, 153995, 109934, 154002, 105516, and 153994. All animal work was approved by the Institutional Animal Care and Use Committee at the University of North Carolina at Chapel Hill under protocols 20-114 and 23-085, following the guidelines outlined by the Association for the Assessment and Accreditation of Laboratory Animal Care and the U.S. Department of Agriculture. All studies were performed in animal biosafety level 3 facilities at the University of North Carolina at Chapel Hill with personnel wearing PAPR, Tyvek suits, Tyvek aprons and booties, and double gloves.

## Acknowledgements

EM data collection was conducted at the Center for Structure Biology Cryo-EM facility at Vanderbilt University. The research was supported by research grants AI167966 and AI158571 to RSB and AI157155 to JEC and RSB. This project was also supported by the Rapidly Emerging Antiviral Drug Development Initiative at the University of North Carolina at Chapel Hill with funding from the North Carolina Coronavirus State and Local Fiscal Recovery Funds program, appropriated by the North Carolina General Assembly. The UNC Animal Histopathology & Laboratory Medicine Core is supported in part by an NCI Center Core Support Grant (5P30CA016086-41) to the UNC Lineberger Comprehensive Cancer Center. We thank Yoshihiro Kawaoka’s lab for the gift of Vero E6 TMPRSS2/ACE2 overexpressing cells. The authors also thank Ryan Lewandowski at Michigan State University for digitally scanning the histologic whole glass slides.

